# Identification of early-stage liver fibrosis by modifications in the interstitial space diffusive microenvironment using fluorescent single-walled carbon nanotubes

**DOI:** 10.1101/2024.02.06.579133

**Authors:** Antony Lee, Apolline A. Simon, Adeline Boyreau, Nathalie Allain-Courtois, Benjamin Lambert, Jean-Philippe Pradère, Fredéric Saltel, Laurent Cognet

**Affiliations:** Laboratoire Photonique Numérique et Nanosciences, Université de Bordeaux, 33400, Talence, France; LP2N, Institut d’Optique Graduate School, CNRS – UMR 5298, 33400, Talence, France; Laboratoire Physico-Chimie Curie, Institut Curie, CNRS - UMR 168, Sorbonne Université 75005, Paris, France; Univ. Bordeaux, CNRS, Bordeaux INP, ICMCB, UMR 5026, 33600 Pessac, France; University of Bordeaux, Inserm, UMR1312, BRIC, Bordeaux Institute of Oncology, 146 Rue Léo Saignat, Bordeaux, F-33076, France; Institut RESTORE - UMR 1301-Inserm/5070-CNRS/EFS/Univ. P. Sabatier, 31037 Toulouse Cedex

**Author notes:** equal contribution (Antony Lee, Apolline A. Simon).

## Abstract

During liver fibrosis, recurrent hepatic injuries lead to accumulation of collagen and other extracellular matrix components in the interstitial space, ultimately disrupting liver functions. Early stages of liver fibrosis may be reversible, but opportunities for diagnosis at these stages are currently limited. Here, we show that the alterations of the interstitial space associated with fibrosis can be probed by tracking individual fluorescent single-walled carbon nanotubes (SWCNTs) diffusing in that space. In a mouse model of early liver fibrosis, we find that nanotubes generally explore elongated areas, whose length decrease as the disease progresses even in regions where histopathological examination does not reveal fibrosis yet. Furthermore, this decrease in nanotube mobility is a purely geometrical effect, as the instantaneous nanotube diffusivity stays unmodified. This work establishes the promise of SWCNTs both for the diagnosis of liver fibrosis at an early stage, and for more in-depth studies of the biophysical effects of the disease.

## Introduction

Liver fibrosis is a pressing public health issue. It is defined as the accumulation of extracellular matrix components, in particular collagen, because of repeated wound healing in response to recurrent injuries, such as viral hepatitis, alcoholic abuse, or nonalcoholic steatohepatitis. This process leads to the accumulation of scar tissue, ultimately leading to cirrhosis, where healthy liver architecture and function are gravely disrupted. While liver fibrosis may be reversible, its onset is typically asymptomatic, which limits the opportunities for early diagnosis, even though the slow progression to cirrhosis, which can take over 15 years, may offer ample time for treatment. While noninvasive tests exist, such as transient elastography (FibroScan) and serum tests, the current reference standard for diagnosis remains the histological examination of liver biopsies, including staining of collagen with Masson’s trichrome or Sirius red^1^, but such approaches fail to detect early stage development of the fibrosis. Moreover, biopsies only represent a small sample of the liver. The fibrotic accumulation of extracellular matrix (ECM) components modifies the physical and chemical properties of interstitial space microenvironment, in particular the ability of small molecules and particles to diffuse in that space that contribute to cell stress. Developing the ability to detect and quantify such changes at an early stage would be relevant, not only for diagnostic purposes, but also because these parameters may affect e.g. the diffusion of drugs through the tissue, and, at a more fundamental level, to understand how these modifications impact disease progression.

The possibility to image single-walled carbon nanotubes (SWCNTs) fluorescence in biological specimens has opened number of possibilities to investigate physio-pathological states of living tissue. SWCNTs present several assets for such purposes. Two key properties that enable imaging SWCNT in deep tissue and live animals are their strong short-wave infrared (SWIR) fluorescence, which matches the biological transparency window, and their large Stokes shift, which minimizes background autofluorescence^2^. Their high brightness and spectacular photostability further allowed single particles detection beyond 50μm deep into tissue slices^3^. In addition, the versality of chiral structures provides several choices of excitation and emission schemes that can be tuned to the tissue’s optical properties^4^, while the possibility to modulate their fluorescence properties by molecular interactions has also allowed detecting the presence of analytes^5^ which can be markers for functional events or diseased states^6–8^.

In this work, we rely on an additional potentiality of SWCNTs formerly demonstrated in brain tissue^3,9,10^, namely, their ability to be detected at the single particle level while they move in the interstitial spaces of live tissue, and thus to access local diffusional environments. This is due to the combination of a nanometric cross-section, which lets the nanotube explore much of the interstitial space without getting stuck, and a micrometric length, which slows down their motion to make fluorescence tracking possible. Based on these assets, we demonstrate here that SWCNT tracking can be tailored to quantify interstitial space characteristics in liver tissue and detect modifications in an early-stage fibrosis model (Figure 1). We find that nanotube trajectories recorded in liver slices display specific nanoscale characteristics that can be analyzed and interpreted as the hallmark of the establishment and development of the disease at very early stage, that remained invisible with current standard diagnosis approaches.

**Figure 1:**
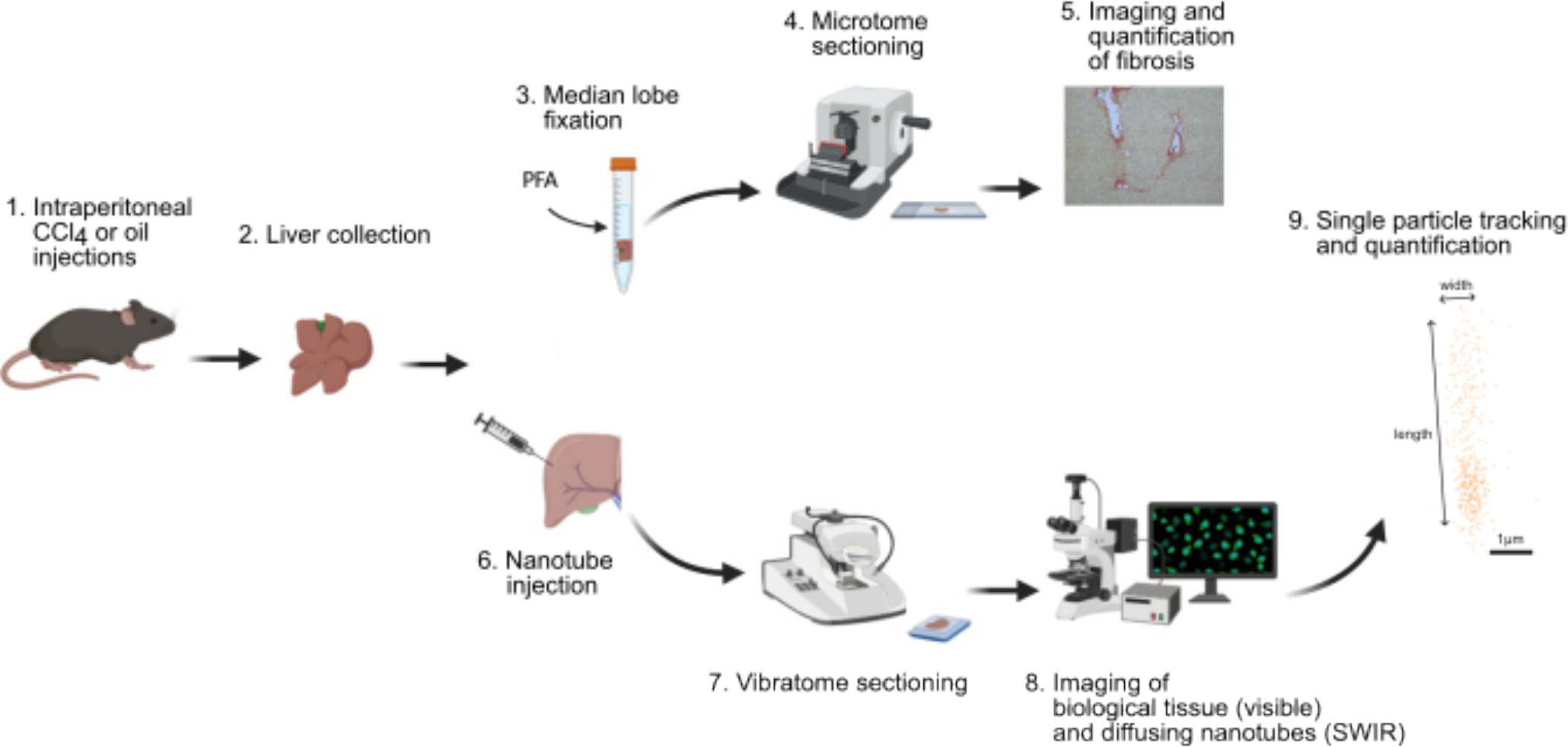
Summary of the ex-vivo experiments for establishing the use of single nanotube tracking in studying early stage liver fibrosis. (1) Mice are injected with CCl_4_ to induce fibrosis (or with oil for the controls), and (2) their livers are collected. On the one hand, (3) the median lobes are fixed, (4) sectioned, and (5) imaged for standard histopathological examination. On the other hand, (6) SWCNTs are injected in the left lobe, which is then (7) sectioned, stained, and (8) imaged; (9) trajectories of nanotubes are reconstructed and quantified.

## Results

### Toxicity & localization of nanotubes in hepatic cell cultures

We first wished to confirm the absence of toxicity and of internalization in hepatic cells of the nanoprobes (18:0 PEG5000 PE-coated nanotubes) that will be used in this study. For this, an in vitro 2D cell model (cultured HuH7 cells, an immortal hepatoma cell line widely used as model in liver research^15,16^), was chosen. We quantified cell proliferation after 3h of growth in presence of 5μg/ml nanotubes in the complete medium, followed by rinsing to eliminate the nanotubes, and cell counting the following day. This protocol is adapted from a previous study of nanotube toxicity on a different human cell line (COS-7)^11^. The proliferated cell number after incubation with nanotubes was 90% that of the cell number in the control condition (without nanotubes). Thus, we conclude that the nanotubes induce no significant toxicity at the concentrations used over a 3h incubation (SI figure 2).

To further test whether 18:0 PEG5000 PE-coated nanotubes would interact with HuH7 cell cultures or even be internalized by the cells, we repeated the above procedure but growing the cells on imaging coverslips. Imaging the cells (including membrane and nuclear staining) immediately after rinsing showed that no nanotubes were left in the sample (SI figure 3A). As a control, we repeated the experiment with PMHC18-mPEG-coated nanotubes, which we found, on the contrary, to be clearly visible in SWIR fluorescence stuck on the cell membranes (as shown by colocalization with the membrane staining) after 3h incubation and rinsing (SI figure 3B). Overall, these results confirm that 18:0 PEG5000 PE-coated nanotubes do not interact nor cross the membrane of HuH7 cells, and can thus be used to study diffusion in the interstitial space of liver tissue. These results are consistent with previous observations that 18:0 PEG5000 PE-coated nanotubes are not internalized in brain tissue cultures^3^, and confirming the capacity of PEG at this molecular weight to function as a biological barrier also observed elsewhere^17^.

### Histopathology

Repeated injections of carbon tetrachloride (CCl_4_) is a commonly used method to induce fibrosis in mice^12^. In this experiment, the apparition and progression of CCl_4_-induced fibrosis was verified by histopathological examination of formaldehyde-fixed median lobe liver slices, performed by Sirius red staining of collagen (figure 2(A, B))^18^. Blinded quantification of the Sirius red staining (figure 2(C)) and grading of the degree of fibrosis was under the Metavir scoring system^13^ confirmed the lack of fibrosis (Metavir F0: no fibrosis) in the control group, the onset of fibrosis (Metavir F1: fibrous expansion of some portal areas ± short fibrous septa) for 4x group, and moderate fibrosis (Metavir F2: fibrous expansion of most portal areas with occasional portal to portal bridging) for 8x group. Importantly, in all these conditions, fibrosis was only detected in the immediate vicinity of the portal areas, but remained invisible in distal regions, which points to the limited sensitivity of the histopathological approach.

**Figure 2:**
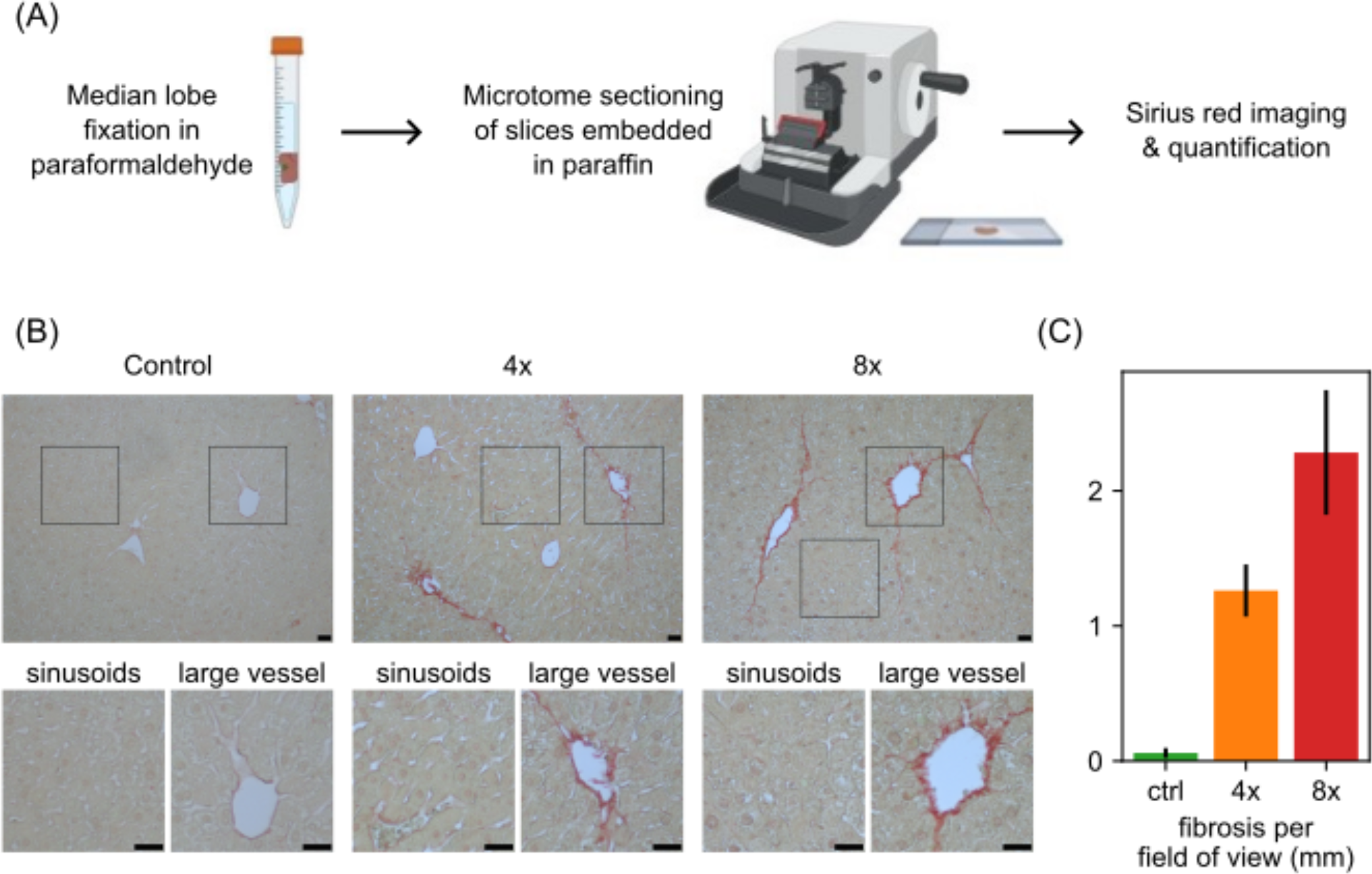
Histological examination of CCl_4_-induced liver fibrosis. (A) Protocol: Liver median lobes are fixed in paraformaldehyde, embedded in paraffin, and sectioned on a microtome, followed by Sirius red staining of collagen fibers and imaging. (B) Sample field of views in control (left), 4x CCl_4_ (middle), and 8x CCl_4_ (right) liver median lobes show the progressive apparition of Sirius-red stained collagen fibers. Insets on each field of view show sinusoids and large vessels; histological examination only shows fibrosis in the latter, not the former. Scale bars, 25μm. (C) Quantification of the average total amount of collagen fibers per field of view also shows the progression of fibrosis.

### Distribution of nanotubes in liver tissue

We next established a protocol for the study of 18:0 PEG5000 PE-coated SWCNT mobility in *ex vivo* liver tissue. It is based on direct nanotube injection into the left lobe followed by vibratome sectioning, then imaging within 15min post-injection. This approach was based on the assumption that the high blood irrigation of liver tissue would ensure fast distribution of the nanotubes throughout the entire tissue. Indeed, we find that this protocol is sufficient to ensure widespread distribution of the nanotubes throughout the tissue, allowing us to easily find field of views where nanotubes were mobile, even far (>1cm) away from the injection site (both for healthy and for diseased tissue).

Fluorescence imaging of liver tissue is intrinsically challenging due to the high autofluorescence generated by the strong blood irrigation. In particular, we found that the previously developed 845 nm excitation and 900nm long-pass emission scheme of (6,5) SWIR nanotubes, determined to be optimal for brain studies^3,19^, was not viable in liver tissue: under these conditions, this tissue exhibits significant autofluorescence, which impedes the ability to track single (6,5) nanotubes (SI figure 4(A)). In this context, we determined that the large Stokes shift of the (7,6) nanotubes (660nm excitation, 1130nm emission) was key in our experiments, as it allows working with a 1100nm long-pass filter that eliminates the significant liver tissue autofluorescence, which occurs at a shorter wavelength (SI figure 4(B)). This new excitation/emission scheme allowed the detection of single SWCNTs *in situ*. In these conditions, 28 nanotubes, on average, were observed per field of view (271μm x 217μm).

Based on this approach, we observed a widespread spatial distribution of the nanotubes, allowing us to avoid imaging in the immediate vicinity injection site, where the tissue structure could have been disrupted by the injection. Multi-color images of the nanotubes and of cellular (nuclear and membrane) stains (figure 3) further allow to identify distal regions from portal areas, where cellular organization appears homogeneous indistinctly of fibrosis progression in the different conditions, a point further confirmed by histopathologic examination. In addition, multi-color images indicate that nanotubes are not internalized and remain in the interstitial space.

**Figure 3:**
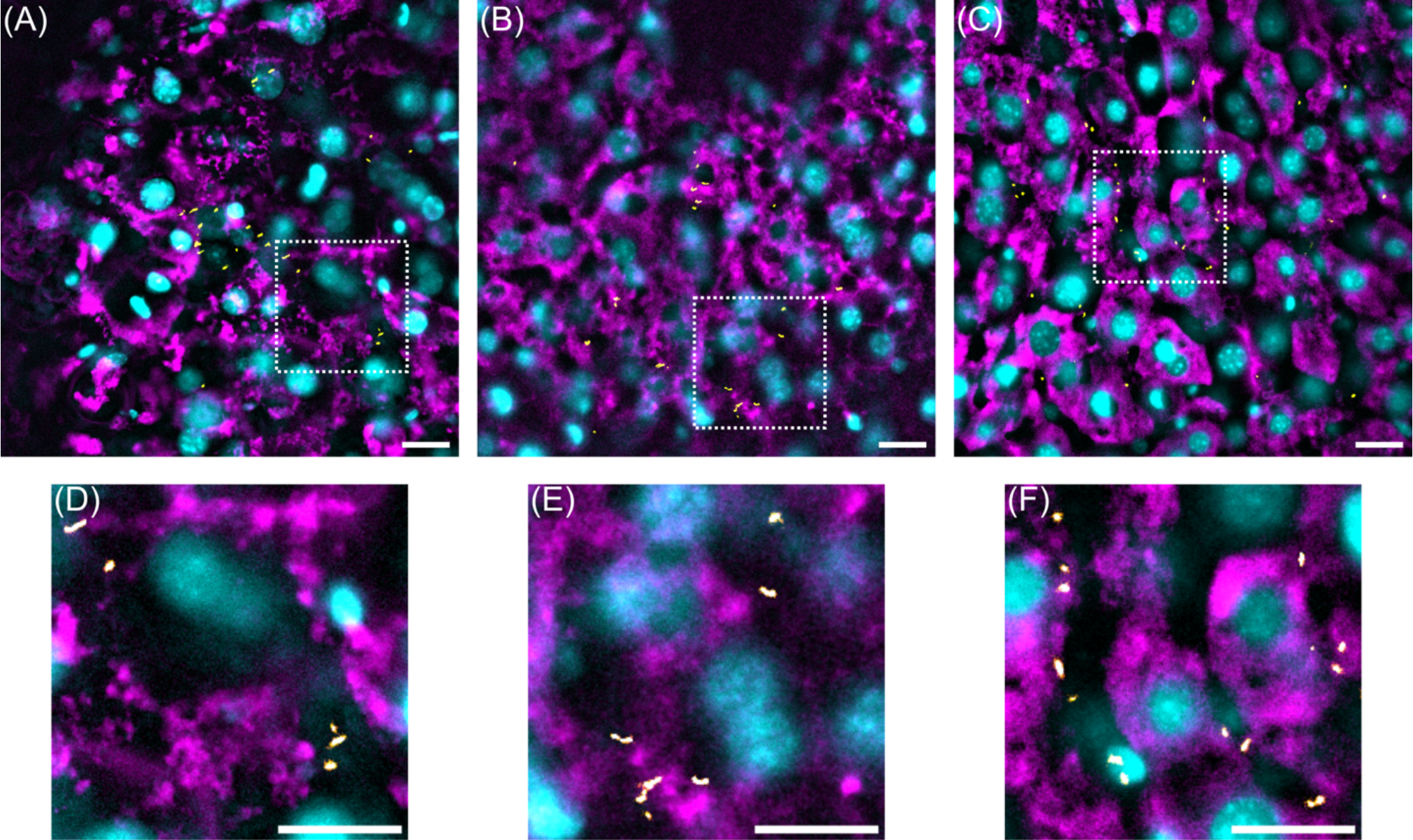
Combined images of cell nuclei (Hoechst staining, cyan), cell membranes (MemGlow staining, magenta), and nanotube trajectories (yellow) in (A) control, (B) 4x CCl_4_, and (C) 8x CCl_4_ mice show that the nanotubes localize to the hepatic interstitial space and exhibit limited mobility. (D), (E), (F); zooms of (A), (B), and (C), respectively, showing the interstitial localization of the nanotubes. Scale bars, 20μm.

Interestingly, all the nanotubes we observed under the microscope showed limited mobility, exploring a region at most a few microns in size. Thus, while the nanotubes are able to diffuse quickly immediately following the injection, they then get quickly trapped into regions of low diffusivity.

### Identification of the fibrosis-induced modifications of the interstitial space dimensions

We observed that the shapes of the areas explored by nanotubes were always relatively featureless ellipses (or disks) (figure 4(A)). Restricting ourselves to nanotubes that were detected over at least 100 frames (which amount to 619 trajectories in the control group, 784 in the 4x CCl_4_ group and 1981 in the 8x CCl_4_ group), we quantified the space explored by the nanotubes by computing the short (width *w*) and the long (length *l*) axis of these ellipses (more specifically, the square roots of the principal inertia moments of the distribution of localizations) (figure 4(A); also summarized in SI figure 5).

**Figure 4:**
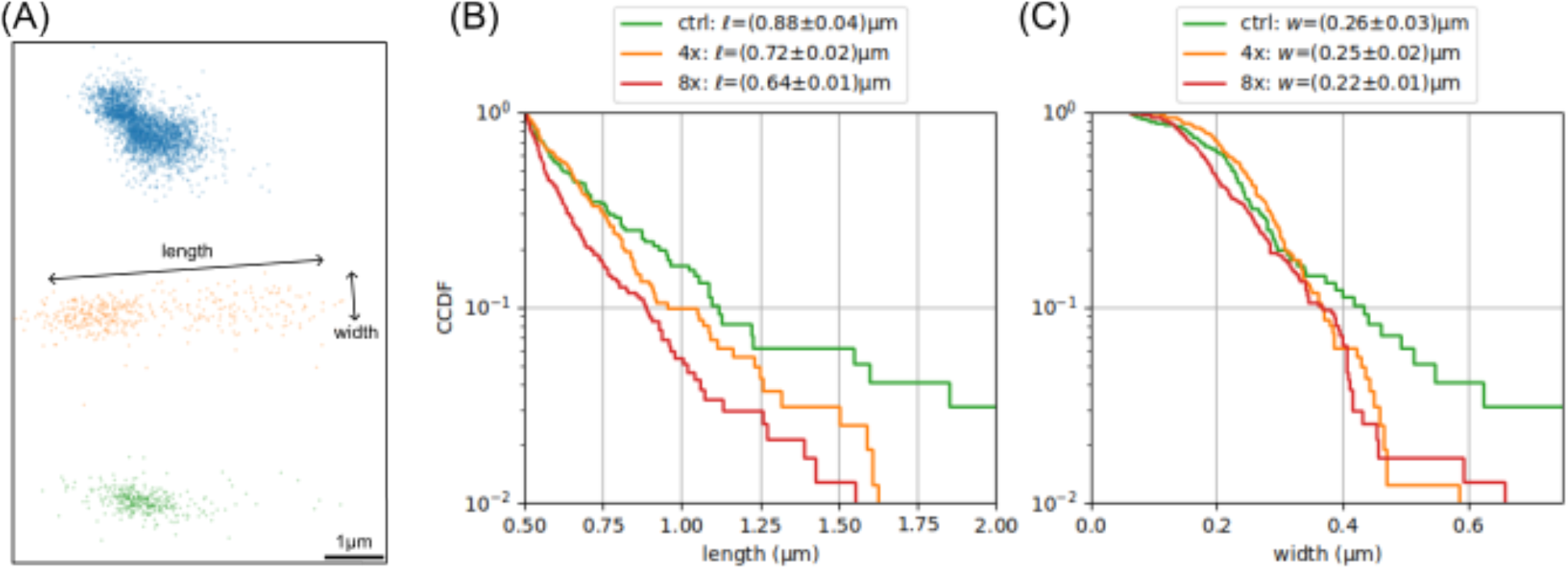
Quantification of the shape of the areas explored by the nanotubes. (A) Set of localizations observed for three sample nanotubes from the control dataset. The area is quasi-elliptical, with a well-defined length *l* and width *w*. (B) The distributions of area lengths *l* for mobile nanotubes (*l*>0.5 μm) of the control, 4x CCl_4_, and 8x CCl_4_ groups show a decrease in mobility as fibrosis progresses. (C) The distributions of area widths *w* for the same nanotubes depend more weakly on the progression of fibrosis; however, both the 4x CCl_4_, and the 8x CCl_4_ groups show a suppression of the greatest widths (>0.5μm).

We refer to nanotubes for which the ellipse long axis is less than 500nm (around half the median nanotube length – 890nm, measured by AFM (SI figure 1(B,C))) as “immobile”; these represent respectively 84%, 79%, and 88% of the control, 4x CCl_4_, and 8x CCl_4_ conditions. The areas explored by these “immobile” nanotubes barely changes between the three conditions: the ellipse length is around (0.34±0.01)μm and the ellipse width around (0.20±0.01)μm (SI figure 6). This latter value also indicates that we can localize single immobile nanotubes over hundreds of frames with an accuracy better than 0.20μm, and thus provides an upper (worst-case) bound on the global accuracy, including both the effects of photon budget and of tissue and microscope drift, achieved in these experiments. We do not consider that the difference across conditions in the fraction of mobile nanotubes is significant either, because these values will also depend on the exact choice of field of views.

On the other hand, the “mobile” nanotubes (*l*>0.5 μm) exhibited significant differences in explored length between the three conditions. In all cases the complementary cumulative distribution function (CCDF) of mobile lengths is approximately exponential (Figure 4(A)), and the mean length (± s.e.) decreases from (0.88±0.04)μm in the control group to (0.72±0.03)μm in the 4x CCl_4_ group and further to (0.64±0.01)μm in the 8x CCl_4_ group. On the other hand, the mobile widths does not change significantly between the three conditions (resp. (0.26±0.03)μm, (0.25±0.02)μm, and (0.22±0.01)μm), although the control group shows more nanotubes with a high mobile width (*w*>0.5μm: 6% of the nanotubes in the control group, 2% in the 4x CCl_4_ group, and 1% in the 8x CCl_4_ group).

Overall, our observations indicate that the explored ellipse length distribution is sensitive to interstitial matrix changes associated with intermediate stages of the disease, whereas the explored ellipse width is not modified.

### Nanotube motion dynamics in the interstitial space

To further quantify the dynamics of nanotube exploration in the interstitial space, we computed the mean-square displacements (MSDs) from the recorded trajectories (Figure 5(A)),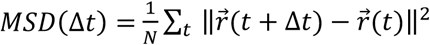. The mean-square displacements display various behaviors, ranging from lack of mobility (constant MSD independent of the time lag Δ*t*) to near-normal diffusion (MSD proportional to the time lag Δ*t*). Nanotubes that exhibit a flat MSD correspond to those defined as “immobile” (*l*<0.5μm) in the previous section. For mobile (*l*>0.5μm) nanotubes, an instantaneous diffusion constant, *D*, is the quarter of the linear rate of increase of the MSD fitted over the first few time lags. We find that the diffusion constant of the mobile nanotubes are approximately exponentially distributed in each condition (figure 5(B)), with negligible change across the conditions (control: *D*=(0.53±0.06)μm2/s, 4x CCl_4_ *D*=(0.59±0.05)μm2/s, 8x CCl_4_ *D*=(0.54±0.04)μm2/s, mean±s.e.). These values can be compared to the expected diffusion constants for micron-sized nanotubes diffusing in water, *D*=2.3μm2/s ^14^; in other words, even though the nanotubes are in an environment whose geometry highly restricts the spatial extent of their motion, the viscosity of the medium is only around 4 times greater than that of water. Conversely, nanotubes internalized in cells have been shown to display effective diffusion coefficients at least 50 times lower than in water^20,21^, which further supports the observation that our nanotubes are not internalized.

**Figure 5:**
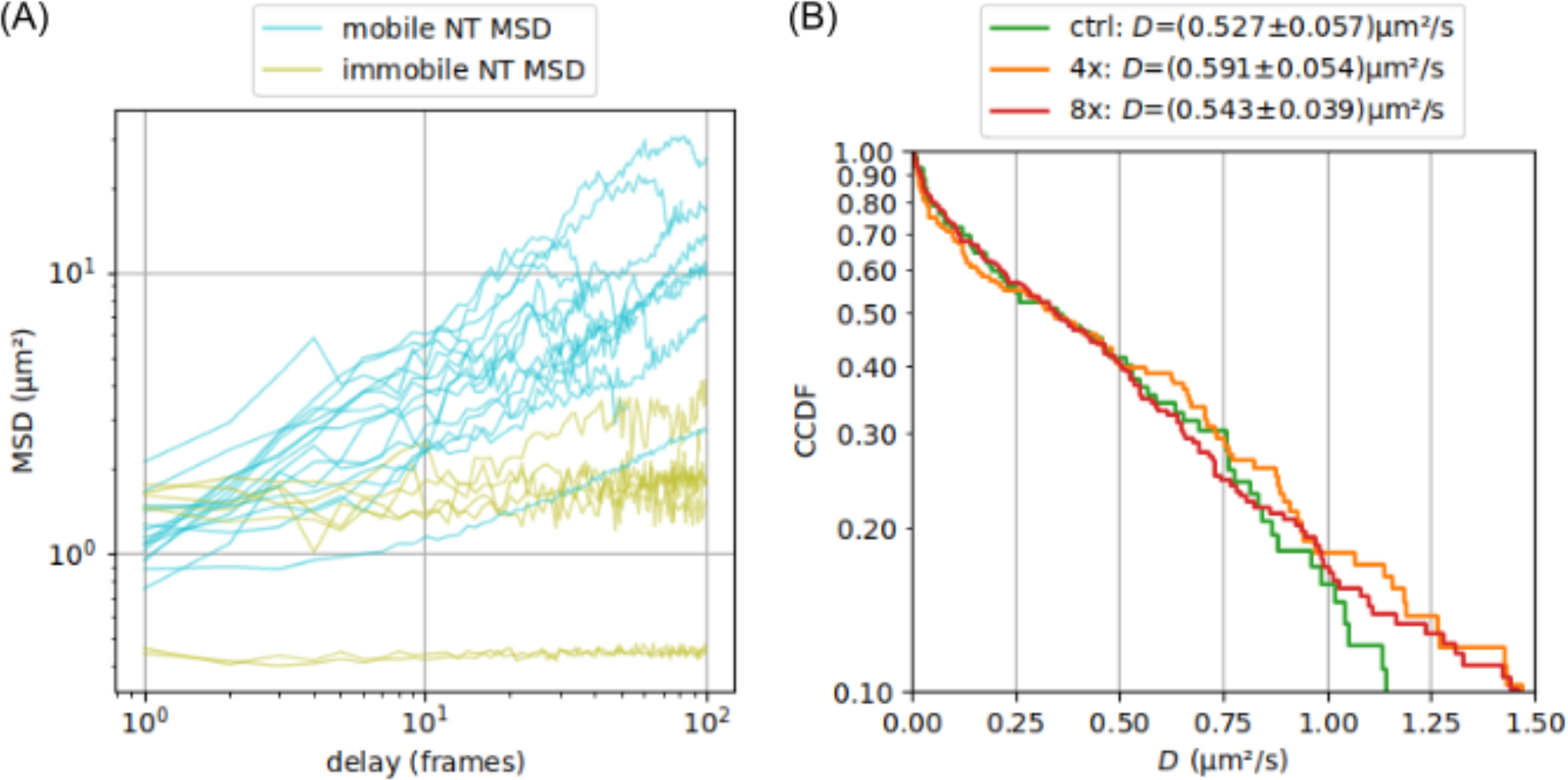
Nanotube mobility analysis. (A) Delay-dependent mean-square displacements of mobile (cyan) and immobile (yellow) nanotubes. (B) The distributions of short-time linear diffusion coefficients of the mobile nanotubes are indistinguishable between the control, 4x CCl_4_, and 8x CCl_4_ conditions.

Overall, this result indicates that the progression of induced fibrosis only affects the instantaneous mobility of the nanotubes in a limited fashion. In particular, the interaction between the phospholipid-PEG coating of the nanotubes and the extracellular matrix is not modified.

## Discussion

The progression of liver fibrosis is associated with modifications of the hepatic extracellular matrix. In this work, we asked whether these changes could be observed at an early stage of the disease by tracking and quantifying the diffusion of fluorescent nanoparticles. We introduce here the use of single-walled carbon nanotubes that bear the advantage of high photostability, excellent tissue penetration due to their thin diameter associated to their 1D character, and wavelength emission versality in the SWIR where tissue exhibit transparency bands. In liver tissue, we used the (7,6) nanotube chirality (660nm excitation, 1130nm emission), to avoid the significant autofluorescence of liver tissue that prevented the use of the (6, 5) chirality (845nm excitation, 985nm emission) (SI figure 3) previously established as an excellent choice in brain tissue^3^. Working on a mouse model of mild liver fibrosis, we established a new protocol to introduce such nanotubes in the hepatic interstitial space and to image them. We observed that nanotubes initially spread quickly and efficiently through the liver tissue, but later show limited mobility in the interstitial space, with a majority of them moving by much less than 0.5μm (half the length of a nanotube), due to being trapped in dead ends of the tortuous interstitial space. Yet, the motion of the diffusing nanotubes provides insights on the modifications of the extracellular matrix induced by fibrosis. We find that in early stages of the disease, in regions away from the portal area and thus not labeled by Sirius red staining, the space that the nanotube can explore is already restricted. The decrease of available space reaches around -18% at the 4x CCl_4_ stage, and a further -9% at the 8x CCl_4_ stage along the long direction of the space explored, but is minor in the transverse direction. We primarily attribute this anisotropy to the asymmetric morphology of the nanotubes, which are around 900nm long but only a few nm in diameter; effectively, nanotubes are confined in “local tubes”, whose length is being more easily probed. Remarkably, this decrease appears to arise purely from the change in the geometry of the local tubes, as the diffusion coefficient of the (mobile) nanotubes does not appear to change significantly; it would be interesting to study whether this parameter changes in later stages of the disease. Overall, these results demonstrate the usefulness of SWCNT tracking to reveal early stage of fibrosis progression in liver tissue, yielding novel insights of changes in the properties of the extracellular matrix, in particular by disentangling the contributions of interstitial space geometry and biochemistry to particle mobility.

## Acknowledgments

This work was performed with financial support from the region Nouvelle-Aquitaine (grant #AAPR2020l-2019-8486010) to L.C and F.S., from CNRS, the France-BioImaging National Infrastructure (ANR-10-INBS-04-01), the Idex Bordeaux (Grand Research Program GPR LIGHT) and the EUR Light S&T (PIA3 Program, ANR-17-EURE-0027) to L.C. A. L. acknowledges support from the Fondation ARC pour la recherche sur le cancer. The authors acknowledge the help of Nathalie Dugot-Senant for the histological studies, performed at the Histopathology Platform, INSERM US 005-CNRS UAR3427 - TBM CORE, a service unit of the CNRS-INSERM and Bordeaux University.

## Materials and methods

### SWIR probe preparation

Biocompatible SWCNT preparation was performed as previously described^11^. Briefly, HiPco nanotubes (Rice University batch 195.7) were suspended in milliQ water in the presence of 5 mg/ml 18:0 PEG5000 PE (Laysan Bio), by homogenization on an Ultra Turrax (IKA T 10 B) at 8000rpm for 10min, probe sonication (Microson, Misonix) for 10min (10W power) on ice, and centrifugation at 20820g for 30 min at 15°C (centrifuge 5804 R, Eppendorf). The supernatant is then kept at 3°C for further use. This suspension was characterized by the measurement of the absorbance spectrum (Evolution 220, Thermo Scientific) and of the luminescence map (SI figure 1(A)). In particular the fluorescence peak at 650nm excitation, 1130nm emission, corresponding to the (7,6) nanotubes chirality, used in this study, is well visible. The nanotube lengths were measured by AFM (SI figure 1(B,C)); the distribution of lengths is centered at (890±470)nm (median±interquartile range).

### Animals and treatments

All experimental procedures were performed in accordance with institutional guidelines for animal research and were approved by the Animal Care and Use Ethics Committee US006 CREFRE-CEEA-

122 (protocol 21-3U1301-JPP-01). All mice were housed under controlled temperature (22–23 °C) and on a 12 h light and 12 h dark cycle with food and water ad libitum.

Hepatic fibrosis was induced in 16-week-old age-matched male C57BL/6 mice (Charles River) by 4 or 8 intraperitoneal injections of carbon tetrachloride (CCl_4_) (0.5 μL/g body weight, diluted 1:3 in corn oil, injected every 3 days). The mice were randomly divided into 3 groups (n = 5 in each group) as follows: control group, with intraperitoneal oil injection and CCl_4_-treated model groups 4 or 8 times^12^.

In all cases, the animals were sacrificed 48hr after the last CCl_4_ injection (in order to observe the fibrotic response rather than the inflammatory alterations that occur immediately post-injection). The liver was extracted from the abdominal cavity using an aseptic technique and placed in a sterile saline solution (Physiodose, Gilbert). The left lobe was immediately prepared an imaged.

### Preparation of liver slices

The protocol for sample preparation is adapted from those previously developed in our lab for brain studies. Around 6.5μg of nanotubes (500μl at 13μg/ml) were manually injected in the left lobe with a 0.6x25mm sterile needle (Agani, Terumo). The lobe was then encased in a block of low-melting point agarose (Thermo Scientific) gel and cut into 800μm-thick slices with a vibratome (Leica VT1200S). The slices farthest from the injection point were placed in an home-made polydimethoxysilane mold on a glass slide, and covered with a solution of phosphate buffer saline (Pan Biotech) with 1‰ Hoechst 33342 (Fisher Scientific) and 2‰ MemGlow 560 (Fluorogenic Membrane Probe, Cystoskeleton) staining, maintained at 28°C by an incubator system, and imaged for up to two hours.

### Imaging

Imaging was performed on an upright Nikon (Eclipse Ni-E) microscope with a 25×, NA 1.10 water immersion objective. Imaging of nanotubes was performed under 660nm laser excitation (OBIS, Coherent), via a 900nm dichroic (FF875-Di01, Semrock) and a 1100nm long-pass filter (FELH1100, Thorlabs), on an InGaAs camera (C-RED2, First Light Imaging). Movies of 4000 frames, at 25 frames per second, were acquired over several field of views per sample.

For each field of view where nanotube motion was observed, imaging of the Hoechst (nuclear) and MemGlow560 (membrane) fluorescence channels was also performed. Hoechst fluorescence images were acquired on an EMCCD camera (ProEM, Princeton Instruments) under mercury lamp illumination via a 387nm excitation filter, a 409nm dichroic, and a 447nm emission filter (387/11, FF409-Di03, and FF02-447/60, Semrock). MemGlow560 fluorescence images were acquired on a sCMOS (Prime BSI, Photometrics) camera under 568nm laser excitation (Sapphire, Coherent) via a 560nm excitation filter, a 580nm dichroic, and a 609nm emission filter (FF01-560/25-25, FF580-Di01 25x36 and FF01-609/62-25, Semrock). This channel was imaged using a HiLo module (Bliq Photonics), a wide-field, speckled-based fluorescence microscopy method which provides confocal-like optical section capabilities. In both cases, a z-stack ranging from - 19μm to +19μm around the nanotube motion plane, by steps of 2μm, was collected.

### Histopathology

The liver median lobe from each mouse was fixed in 4% formaldehyde (Fisher Scientific) and embedded in paraffin for later histological analysis. Each sample was cut into 5 μm-thick sections and stained with Sirius red (VWR) according to standard procedures for identifying collagen deposition. Bright-field imaging of the labelled sections (8 fields of view per mouse) was performed (LEICA DMR microscope coupled with a NIKON DS-FI2 camera) ; the stained collagen fibers was then manually quantified (NIS BR imaging software) and graded following the Metavir scoring system^13^ in a blinded manner by a single pathologist.

### Data processing

Nanotube movies were analyzed using custom Python and C++ programs that performed detected, super-localized, and tracked the fluorescent particles to reconstruct their trajectories^14^. Only trajectories with at least 20 localizations were kept for further analysis. A total of 4079 trajectories, corresponding to 3.7x10^6^ localizations over 145 field of views (271μm x 217μm), were reconstructed.

### Cell culture

Human hepatoma cell lines HuH7 (ATCC)^15^ used for all in vitro experiment were grown in high-glucose Dulbecco’s modified Eagle’s medium with L-Glutamine and sodium pyruvate (Pan Biotech, P03-0810) supplemented with 1% of penicillin 10000U and streptomycin (10mg/ml), and 10% fetal bovine serum (FBS-16A CliniSciences). Cells were subcultured twice a week in T75 flasks until 80% confluent, then trypsinized, centrifuged, and resuspended with a 1:5 split ratio in new T75.

For cytotoxicity and localization tests, 10^5^ HuH7 cells were plated per well in sterile twelve-well plates (growth area 3.8 cm^2^, CLEARLine), allowed to adhere overnight to 80% confluence, and then exposed to culture media alone or media containing 5µg/ml nanotubes during 3h. Each condition was duplicated. Cell proliferation and toxicity were determined by direct counting cell suspension of each well with Neubauer counting chambers (Marienfeld Superior 0650030). Counting and cell morphology verification were performed by bright field imaging on an inverted microscope (Nikon Eclipse Ts2, Nikon camera, 4x or 10x magnification).

To establish where nanotubes would localize in a liver cell culture, HuH7 cells were grown on glass coverslips in a petri dish. Once the cells reached 80% confluence, they were exposed to media containing 5μg/ml nanotubes (coated either with PEG5000 PE or with PMHC18-mPEG (control)) for 3hr, then the coverslip was washed and transferred to a Ludin chamber (Life Imaging Services), stained with 1‰ Hoechst 33342 (Fisher Scientific) and 2‰ MemGlow 560 (Fluorogenic Membrane Probe, Cytoskeleton), and imaged on the imaging setup described below.

## References

(1) Ginès, P.; Castera, L.; Lammert, F.; Graupera, I.; Serra-Burriel, M.; Allen, A. M.; Wong, V. W.; Hartmann, P.; Thiele, M.; Caballeria, L.; De Knegt, R. J.; Grgurevic, I.; Augustin, S.; Tsochatzis, E. A.; Schattenberg, J. M.; Guha, I. N.; Martini, A.; Morillas, R. M.; Garcia-Retortillo, M.; De Koning, H. J.; Fabrellas, N.; Pich, J.; Ma, A. T.; Diaz, M. A.; Roulot, D.; Newsome, P. N.; Manns, M.; Kamath, P. S.; Krag, A.; for the LiverScreen Consortium Investigators. Population Screening for Liver Fibrosis: Toward Early Diagnosis and Intervention for Chronic Liver Diseases. Hepatology 2022, 75 (1), 219–228. 10.1002/hep.32163.

(2) Hong, G.; Diao, S.; Antaris, A. L.; Dai, H. Carbon Nanomaterials for Biological Imaging and Nanomedicinal Therapy. Chem. Rev. 2015, 115 (19), 10816–10906. 10.1021/acs.chemrev.5b00008.

(3) Godin, A. G.; Varela, J. A.; Gao, Z.; Danné, N.; Dupuis, J. P.; Lounis, B.; Groc, L.; Cognet, L. Single-Nanotube Tracking Reveals the Nanoscale Organization of the Extracellular Space in the Live Brain. Nature Nanotech 2017, 12 (3), 238–243. 10.1038/nnano.2016.248.

(4) Weisman, R. B.; Bachilo, S. M. Dependence of Optical Transition Energies on Structure for Single-Walled Carbon Nanotubes in Aqueous Suspension: An Empirical Kataura Plot. Nano Lett. 2003, 3 (9), 1235–1238. 10.1021/nl034428i.

(5) Barone, P. W.; Baik, S.; Heller, D. A.; Strano, M. S. Near-Infrared Optical Sensors Based on Single-Walled Carbon Nanotubes. Nature Mater 2005, 4 (1), 86–92. 10.1038/nmat1276.

(6) Beyene, A. G.; Delevich, K.; Del Bonis-O’Donnell, J. T.; Piekarski, D. J.; Lin, W. C.; Thomas, A. W.; Yang, S. J.; Kosillo, P.; Yang, D.; Prounis, G. S.; Wilbrecht, L.; Landry, M. P. Imaging Striatal Dopamine Release Using a Nongenetically Encoded near Infrared Fluorescent Catecholamine Nanosensor. Sci. Adv. 2019, 5 (7), eaaw3108. 10.1126/sciadv.aaw3108.

(7) Elizarova, S.; Chouaib, A. A.; Shaib, A.; Hill, B.; Mann, F.; Brose, N.; Kruss, S.; Daniel, J. A. A Fluorescent Nanosensor Paint Detects Dopamine Release at Axonal Varicosities with High Spatiotemporal Resolution. Proc. Natl. Acad. Sci. U.S.A. 2022, 119 (22), e2202842119. 10.1073/pnas.2202842119.

(8) Antman-Passig, M.; Yaari, Z.; Goerzen, D.; Parikh, R.; Chatman, S.; Komer, L. E.; Chen, C.; Grabarnik, E.; Mathieu, M.; Haimovitz-Friedman, A.; Heller, D. A. Nanoreporter Identifies Lysosomal Storage Disease Lipid Accumulation Intracranially. Nano Lett. 2023, acs.nanolett.3c02502. 10.1021/acs.nanolett.3c02502.

(9) Soria, F. N.; Paviolo, C.; Doudnikoff, E.; Arotcarena, M.-L.; Lee, A.; Danné, N.; Mandal, A. K.; Gosset, P.; Dehay, B.; Groc, L.; Cognet, L.; Bezard, E. Synucleinopathy Alters Nanoscale Organization and Diffusion in the Brain Extracellular Space through Hyaluronan Remodeling. Nat Commun 2020, 11 (1), 3440. 10.1038/s41467-020-17328-9.

(10) Paviolo, C.; Ferreira, J. S.; Lee, A.; Hunter, D.; Calaresu, I.; Nandi, S.; Groc, L.; Cognet, L. Near-Infrared Carbon Nanotube Tracking Reveals the Nanoscale Extracellular Space around Synapses. Nano Lett. 2022, 22 (17), 6849–6856. 10.1021/acs.nanolett.1c04259.

(11) Gao, Z.; Danné, N.; Godin, A.; Lounis, B.; Cognet, L. Evaluation of Different Single-Walled Carbon Nanotube Surface Coatings for Single-Particle Tracking Applications in Biological Environments. Nanomaterials 2017, 7 (11), 393. 10.3390/nano7110393.

(12) Scholten, D.; Trebicka, J.; Liedtke, C.; Weiskirchen, R. The Carbon Tetrachloride Model in Mice. Lab Anim 2015, 49 (1_suppl), 4–11. 10.1177/0023677215571192.

(13) Goodman, Z. D. Grading and Staging Systems for Inflammation and Fibrosis in Chronic Liver Diseases. Journal of Hepatology 2007, 47 (4), 598–607. 10.1016/j.jhep.2007.07.006.

(14) Lee, A.; Cognet, L. Length Measurement of Single-Walled Carbon Nanotubes from Translational Diffusion and Intensity Fluctuations. Journal of Applied Physics 2020, 128 (22), 224301. 10.1063/5.0031194.

(15) Ezzoukhry, Z.; Henriet, E.; Piquet, L.; Boyé, K.; Bioulac-Sage, P.; Balabaud, C.; Couchy, G.; Zucman-Rossi, J.; Moreau, V.; Saltel, F. TGF-B1 Promotes Linear Invadosome Formation in Hepatocellular Carcinoma Cells, through DDR1 up-Regulation and Collagen I Cross-Linking. European Journal of Cell Biology 2016, 95 (11), 503–512. 10.1016/j.ejcb.2016.09.003.

(16) Kasai, F.; Hirayama, N.; Ozawa, M.; Satoh, M.; Kohara, A. HuH-7 Reference Genome Profile: Complex Karyotype Composed of Massive Loss of Heterozygosity. Human Cell 2018, 31 (3), 261–267. 10.1007/s13577-018-0212-3.

(17) Owens, D.; Peppas, N. Opsonization, Biodistribution, and Pharmacokinetics of Polymeric Nanoparticles. International Journal of Pharmaceutics 2006, 307 (1), 93–102. 10.1016/j.ijpharm.2005.10.010.

(18) Huang, Y.; De Boer, W. B.; Adams, L. A.; MacQuillan, G.; Rossi, E.; Rigby, P.; Raftopoulos, S. C.; Bulsara, M.; Jeffrey, G. P. Image Analysis of Liver Collagen Using Sirius Red Is More Accurate and Correlates Better with Serum Fibrosis Markers than Trichrome. Liver International 2013, 33 (8), 1249–1256. 10.1111/liv.12184.

(19) Danné, N.; Godin, A. G.; Gao, Z.; Varela, J. A.; Groc, L.; Lounis, B.; Cognet, L. Comparative Analysis of Photoluminescence and Upconversion Emission from Individual Carbon Nanotubes for Bioimaging Applications. ACS Photonics 2018, 5 (2), 359–364. 10.1021/acsphotonics.7b01311.

(20) Reuel, N. F.; Dupont, A.; Thouvenin, O.; Lamb, D. C.; Strano, M. S. Three-Dimensional Tracking of Carbon Nanotubes within Living Cells. ACS Nano 2012, 6 (6), 5420–5428. 10.1021/nn301298e.

(21) Fakhri, N.; Wessel, A. D.; Willms, C.; Pasquali, M.; Klopfenstein, D. R.; MacKintosh, F. C.; Schmidt, C. F. High-Resolution Mapping of Intracellular Fluctuations Using Carbon Nanotubes. Science 2014, 344 (6187), 1031–1035. 10.1126/science.1250170.

