## Supplementary Figures 1-6 for "Identification of early-stage liver fibrosis by modifications in the interstitial space diffusive microenvironment using fluorescent single-walled carbon nanotubes"

**equal contribution*

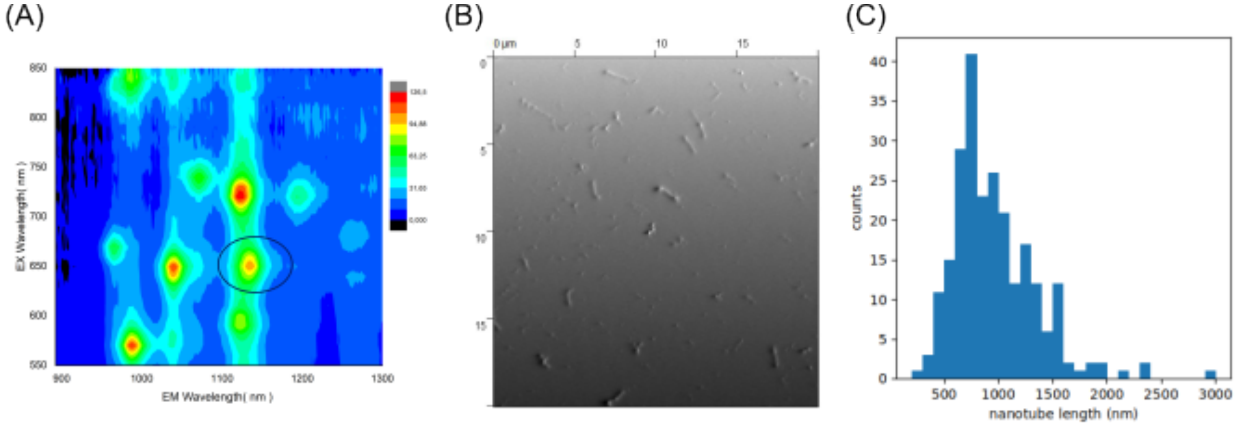


**SI figure 1:** ***Nanotube characterization.*** (A) Photoluminescence excitation map. Acquisition between 400 nm and 850 nm with a 5 nm step for the excitation and between 900 nm and 1400 nm for the emission with also a 5 nm step. The integration time used is 10 seconds. (B) AFM image of nanotubes, used to determine (B) a histogram of the nanotubes length (*N*=241 nanotubes).

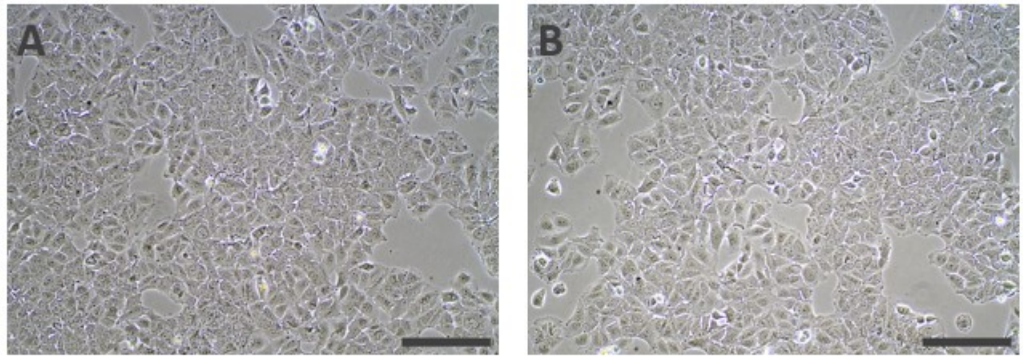


**SI figure 2:** ***Bright field images of HuH7 cells***: (A) control, (B) after 3h of incubation with 8:0 PEG5000 PE-coated nanotubes. No significant difference is observed (see main text) Scale bar: 500 µm.

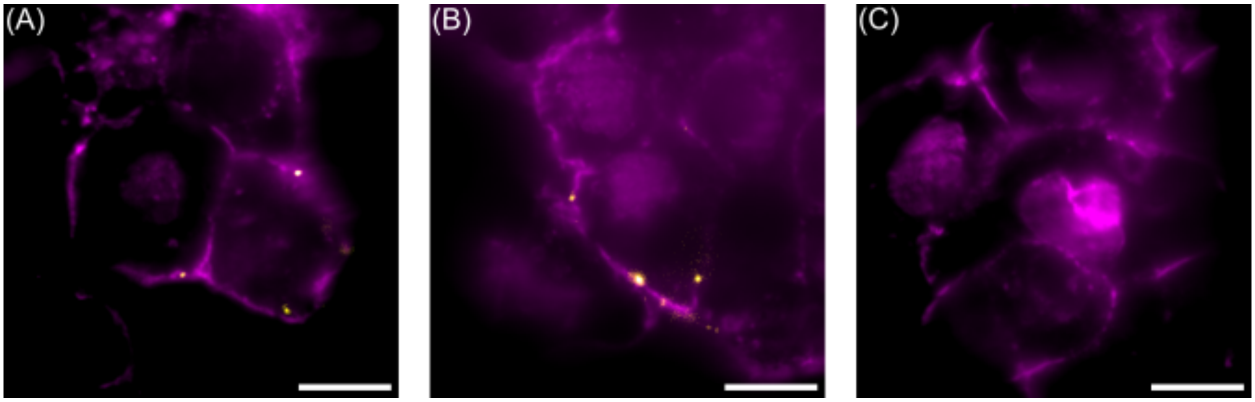


**SI figure 3:** ***Testing for the interaction between nanotubes and cell membranes***. (A, B) PMHC18-mPEG-coated nanotubes (yellow) incubated with HuH7 cells stick to cell membranes (magenta) even after rinsing. (C) 18:0 PEG5000 PE-coated nanotubes incubated with HuH7 cells disappear after rinsing, indicating a fortiori that they are not internalized.

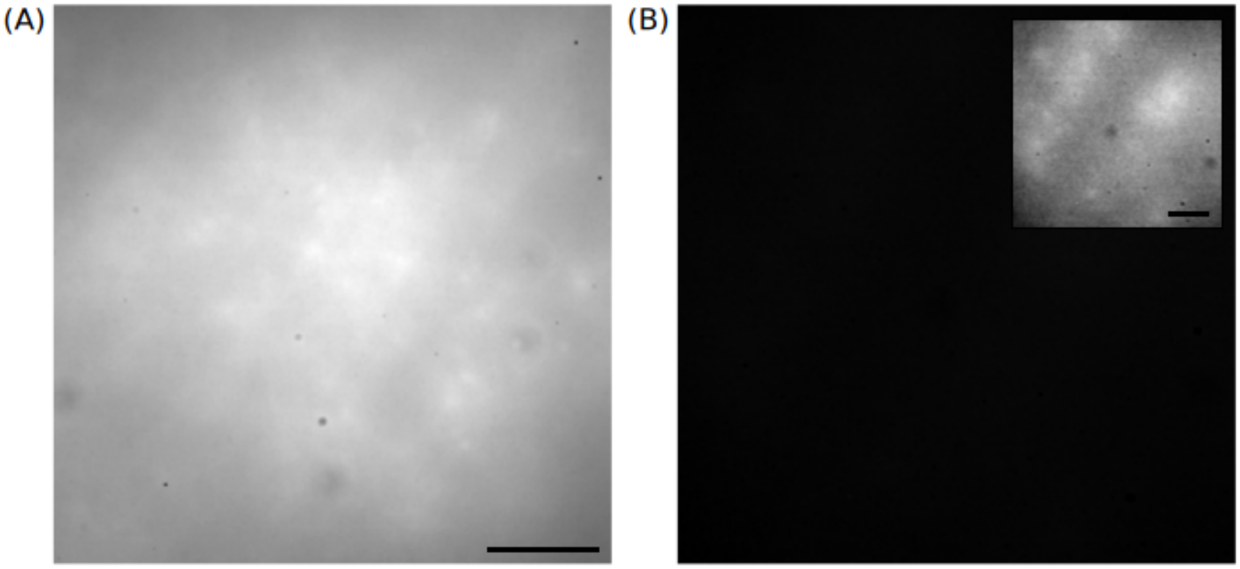


**SI figure 4:** ***Comparison of liver tissue autofluorescence*** under (A) 845nm excitation and a 900nm long-pass emission filter, suitable for imaging (6, 5) nanotubes, and (B) 660nm excitation and a 1100nm long-pass emission filter, suitable for imaging (7, 6) nanotubes. The tissue exhibits significant autofluorescence only under the former imaging conditions, whereas no autofluorescence is visible at all under the latter. (B) inset: Increasing the contrast 20-fold reveals the very weak autofluorescence signal still present under 660nm excitation and 1100nm long-pass emission. Scale bars, 20μm.

| group | count | length (µm) | width (µm) | diffusion const. (µm²/s) |
|---|---|---|---|---|
| ctrl-immobile | 522 (84%) | 0.34±0.01 | 0.20±0.01 | |
| 4x-immobile | 623 (79%) | 0.34±0.01 | 0.21±0.01 | |
| 8x-immobile | 1746 (88%) | 0.33±0.01 | 0.19±0.01 | |
| ctrl-mobile | 97 (16%) | 0.88±0.04 | 0.26±0.03 | 0.53±0.06 |
| 4x-mobile | 161 (21%) | 0.72±0.02 | 0.25±0.02 | 0.59±0.05 |
| 8x-mobile | 235 (12%) | 0.64±0.01 | 0.22±0.01 | 0.54±0.04 |

**SI figure 5:** ***Summary of trajectory characteristics.***

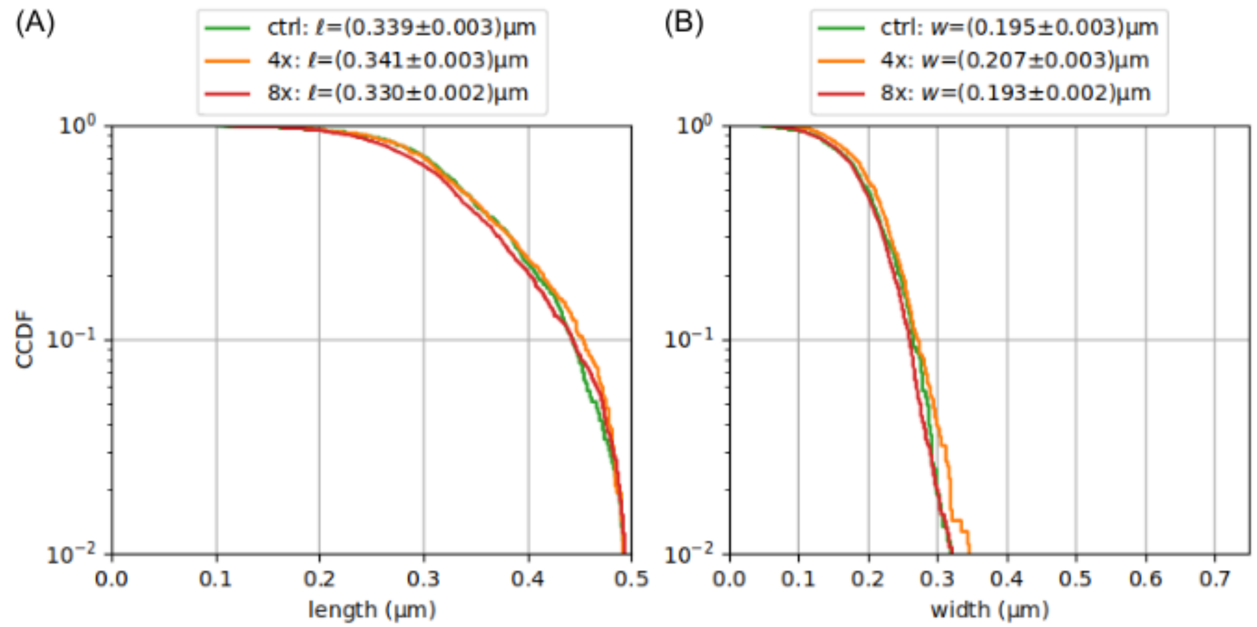


**SI figure 6:** The distributions of area (A) lengths and (B) widths of immobile nanotubes show no variation between the control, 4x $CCl_4$, and 8x $CCl_4$ conditions.
